# Repression of *MRP51* in cis does not contribute to the synthetic growth defect conferred by an *hphMX4*-marked deletion of *DBP1* in a *ded1-ts* mutant

**DOI:** 10.1101/2024.01.30.578007

**Authors:** Fan Zhang, Neelam Dabas Sen, Alan G. Hinnebusch

**Affiliations:** Division of Molecular and Cellular Biology, Eunice K. Shriver National Institute of Child Health and Human Development, National Institutes of Health, Bethesda, Maryland 20892, USA; School of Life Sciences, Jawaharlal Nehru University, New Delhi 110067, India

**Keywords:** Dbp1, Ded1, helicase, translation, initiation, yeast

## Abstract

Powers et al. recently demonstrated that the *hphMX6* cassette used to delete *DPB1* in *dbp1Δ::hphMX6* yeast mutants leads to reduced expression in *cis* of the adjacent gene *MRP51*, encoding the mitochondrial small subunit (SSU) ribosomal protein Mrp51. Here we provide evidence that elimination of Dbp1, not reduced *MRP51* expression, underlies the synthetic growth defect of a *dbp1Δ::hphMX6 ded1-ts* mutant on glucose-containing medium, where respiration is dispensable, consistent with our previous conclusion that Dbp1 and Ded1 perform overlapping functions in stimulating translation initiation on mRNAs burdened with long or structured 5’UTRs in cells cultured with glucose.

## INTRODUCTION

Structures in mRNA that impede ribosome binding or subsequent scanning of the 5’untranslated region (5’UTR) to locate the AUG initiation codon reduce translation efficiency. An abundance of evidence indicates that yeast DEAD-box RNA helicase Ded1 promotes translation initiation by enhancing ribosomal scanning of long or structure-laden 5’UTRs (1-6). Previously, we published ribosome profiling data suggesting that Dbp1, a Ded1 paralog, performs similar functions in yeast cells, finding that many more mRNAs exhibit reduced relative translational efficiencies (TEs) in a *ded1-ts dbp1Δ* double mutant than in either single mutant, with numerous transcripts exhibiting strong dependence on Dbp1 or Ded1 for efficient translation only when the other helicase is impaired in the double mutant. Such “conditionally hyperdependent” mRNAs contain unusually long 5′UTRs with heightened propensities for secondary structure formation—the same characteristics found previously for mRNAs translationally impaired in the *ded1-ts* single mutant (2). Consistently, overexpressing Dbp1 in a *ded1-cs* mutant suppressed the growth defect and improved the TEs of many Ded1-hyperdependent mRNAs in this single mutant. Moreover, we found that Dbp1 associates with translating mRNAs in cell extracts and that purified Dbp1 resembled Ded1 in preferentially accelerating 48S PIC assembly on selected mRNAs harboring structured 5’UTRs in a fully reconstituted yeast translation initiation system. Finally, profiling of 40S initiation complexes in the *ded1* and *dbp1* mutants provided evidence that Ded1 and Dbp1 stimulate PIC attachment or scanning of 5’UTRs on many Ded1/Dbp1 hyperdependent mRNAs in vivo (6).

Subsequently, Powers et al. reported (7) that the *hphMX6* cassette we had utilized in deleting *DBP1* in *dbp1Δ::hphMX6* strains in the aforementioned study (6) leads to reduced expression in *cis* of the adjacent gene *MRP51*, encoding the mitochondrial small subunit (SSU) ribosomal protein Mrp51. This defect was shown to be associated with impaired mitochondrial SSU biogenesis, reduced bulk cytoplasmic translation, and diminished cell growth on media containing non-fermentable carbon sources in which respiration is required for energy generation. These phenotypes could be suppressed by introduction of *MRP51*, but not *DBP1*, on a plasmid. The inhibition of *MRP51* expression appears to involve a transcript emanating from the strong *TEF1* promoter of the *MX6* cassette that overlaps the native *MRP51* promoter and, although encompassing the entire *MRP51* ORF, is likely impaired for Mrp51 production owing to alternative translation initiation events at upstream open-reading-frames in the extended transcript. Presumably, production of this extended transcript interferes with synthesis of the native *MRP51* mRNA in the manner described previously for long-untranslated-transcript-isoforms (LUTIs) in WT yeast cells (8,9). A similar reduction in *MRP51* expression was observed by Powers et al. (7) on interrogating our published ribosome profiling and RNA-Seq data for the *dbp1Δ::hphMX4* mutant grown on glucose-containing medium, resulting in a ∼4-fold decrease in translation of *MRP51* mRNA, from which we had concluded erroneously that Dbp1 enhances the translational efficiency (TE) of *MRP51* mRNA (6).

All of the experiments in our previous analysis of *dbp1Δ::hphMX6* mutants (6) were conducted on cells growing in medium containing glucose as carbon source, where Mrp51 and respiration in mitochondria are not essential for energy generation. Indeed, we showed that the *dbp1Δ::hphMX6* single mutant is indistinguishable from WT both in cell growth rate and bulk polysome assembly (6), indicating that any reduction in mitochondrial translation that results from reduced *MRP51* expression has no discernible impact on bulk cytoplasmic translation in glucose-grown cells. Similar to our findings, De la Cruz et al. analyzed a *dbp1Δ:TRP1* allele and found a synthetic slow growth phenotype (Slg^-^) in combination with a *ded1/spp81-3* mutation on glucose-containing medium, despite the absence of any Slg^-^ for the *dbp1Δ::TRP1* mutation; and no synthetic interaction was observed on combining *dbp1Δ::TRP1* with mutations impairing other initiation factors, including *tif1–1* (eIF4A), *cdc33–1* and *cdc33–42* (eIF4E), Δ*stm1* (eIF4B), or Δ*tif4631* (eIF4G1) (10). We consider it unlikely that a reduction in *MRP51* expression conferred by *dbp1Δ::hphMX4* or *dbp1Δ:TRP1* would produce such a highly specific exacerbation of the phenotypes of *ded1* mutations in glucose-grown cells, especially considering that the *TRP1* promoter is >10 times weaker than the *S. cerevisiae TEF1* promoter on glucose media (11).

Nevertheless, we could not eliminate the possibility that a reduction in *MRP51* expression was contributing to the synthetic reductions in growth conferred by the *dbp1Δ::hphMX4* mutation in the *dbp1Δ::hphMX ded1-ts* double mutant—a key observation indicating functional cooperation between Dbp1 and Ded1 in stimulating translation (6). We had shown that the synthetic Slg^-^ phenotype of the *ded1-ts dbp1Δ::hphMX4* double mutant was fully complemented by *DBP1* on a high-copy plasmid (lacking *MRP51*); however, this is an imperfect experiment because *DBP1* overexpression can suppress the growth defect of *ded1* mutations (as noted above) in addition to complementing the *dbp1Δ* deletion. As such, it was still possible that reduced *MRP51* expression conferred by *dbp1Δ::hphMX4*, rather than loss of *DBP1* function, was responsible for exacerbating the growth defect conferred by the *ded1-ts* mutation in the *ded1-ts dbp1Δ::hphMX4* double mutant on glucose medium. In an effort to rectify this shortcoming, we tested whether a low-copy plasmid containing *DBP1* but lacking *MRP51* could likewise complement the synthetic Slg^-^ phenotype of the *dbp1Δ::hphMX4 ded1-ts* double mutant, whereas a plasmid harboring *MRP51* alone could not.

## RESULTS AND DISCUSSION

As expected, the low-copy *DED1* plasmid fully complemented the strong Slg^-^ of the *dbp1Δ::hphMX4 ded1-ts* double mutant examined in our previous study (6) and conferred WT growth at 34°C on medium with glucose as carbon source (Figure 1A, row 5 vs. rows 4, 1, & 3). These results are consistent with our previous finding that deletion of *DBP1* alone by *dbp1Δ::hphMX4* does not appreciably impair cell growth or bulk translation on glucose-containing medium (6). Importantly, introducing the low-copy *DBP1* plasmid into the double mutant increased growth at 34°C compared to the empty vector control, but left a residual growth defect intact in comparison to the WT strain, comparable to that exhibited by the *ded1-ts* single mutant (row 6 vs. rows 1-2). In contrast, introducing low-copy *MRP51* afforded no growth improvement at 34°C and was indistinguishable from empty vector (row 7 vs. row 4). These results indicate that the stronger growth defect of the *dbp1Δ::hphMX4 ded1-ts* double mutant compared to the *ded1-ts* single mutant results from loss of *DBP1*, not *MRP51*, function.

**Figure 1.**
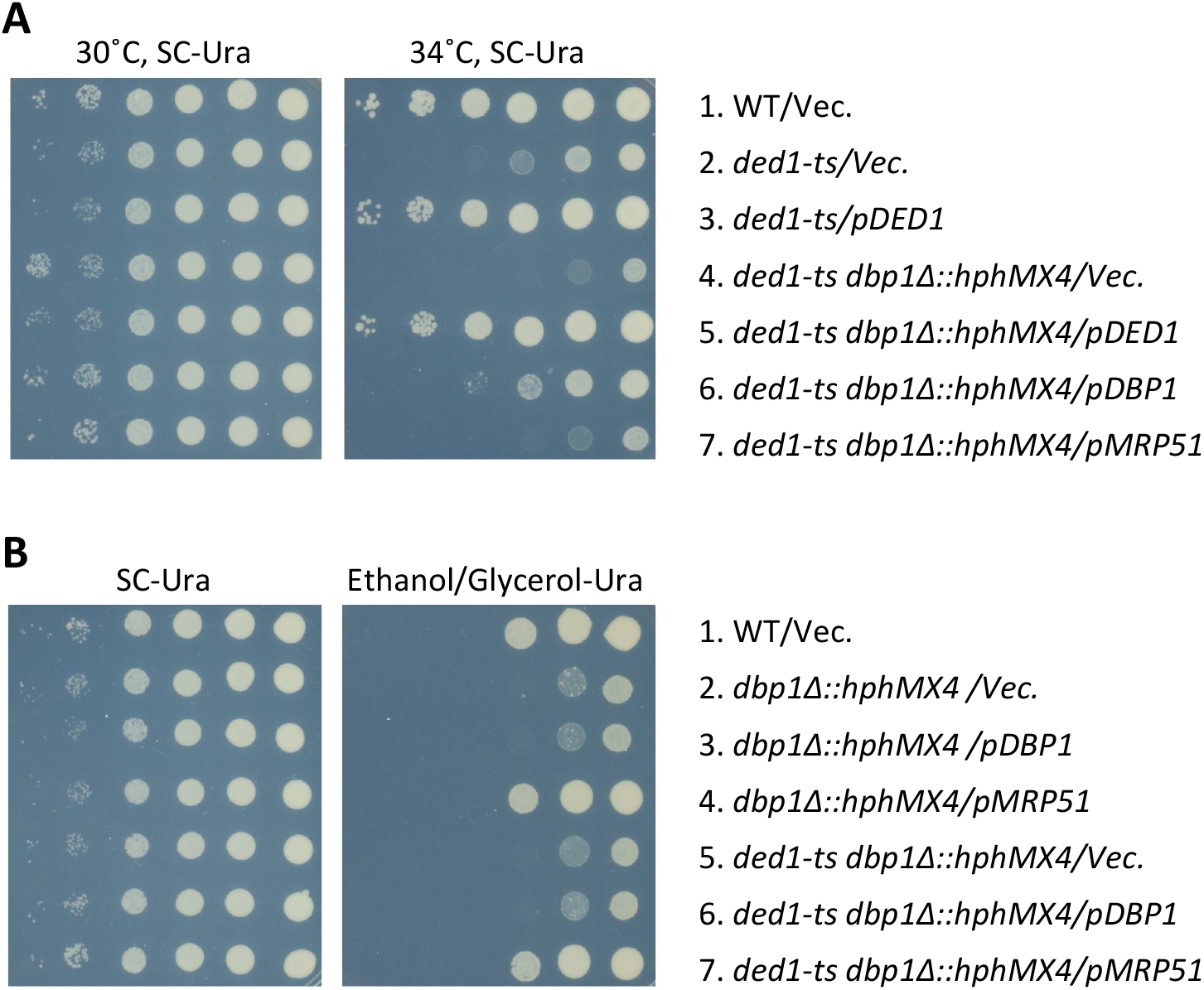
Low-copy *DBP1* but not *MRP51* complements the slow-growth phenotype conferred by *dbp1Δ::hphMX4* in the *ded1-ts dbp1Δ::hphMX4* double mutant on glucose-containing medium. **(A)** Serial dilutions of the following yeast strains were spotted on synthetic complete (SC) medium without uracil containing 2% glucose as carbon source and incubated at 30°C or 34°C: WT strain BY4741 (*MATa his3Δ1 leu2Δ0 met15-Δ0 ura3-Δ0)* transformed with low-copy *URA3 CEN6* vector pRS416 (12) (row 1), strain Y10029 (aka F2030) (*MATa his3Δ1 leu2Δ0 met15-Δ0 ura3-Δ0 ded1-952::kanMX4* (13)) transformed with pRS416 *or* low-copy *URA3* plasmid CP3482 containing *DED1* (14) (rows 2-3), and strain H5331 (aka NSY80) (*MATa his3Δ1 leu2Δ0 met15-Δ0 ura3-Δ0 ded1-952::kanMX4 dbp1Δ::hphMX4* pRS413 [*HIS3*]) transformed with pRS416, CP3482, low-copy *URA3* plasmid pNDS42 (aka p6251) containing *DBP1* (6), or low-copy *URA3* plasmid pRS416-MRP51 (aka p6749) containing *MRP51* (rows 4-7). Strain H5331 is a transformant of NSY15 (6) containing low-copy *HIS3* vector pRS413. **(B)** Serial dilutions of the following strains were spotted on synthetic complete medium lacking uracil and containing 2% glycerol/2% ethanol in place of glucose as carbon source and incubated at 30°C: WT strain BY4741 transformed with pRS416 (row 1), strain H5314 (aka NSY14) (*MATa his3Δ1 leu2Δ0 met15-Δ0 ura3-Δ0 dbp1Δ::hphMX4* (6) transformed with pRS416, pNDS42, or p6749 (rows 2-4), and H5331 transformed with pRS416, pNDS42, or p6749 (rows 5-7). Plasmid p6749 was constructed by synthesis of a Bam HI-Hind III DNA fragment containing the 1035 bp of the *MRP51* ORF, 360 bp 5’ of the ORF, and 200 bp 3’ of the ORF (carried out by LifeSct LLC) and inserted between the Bam HI and Hind III sites of pRS416. The cloned synthesized fragment was sequenced in its entirety to verify the WT sequence.

In sharp contrast to these last results, on medium containing the non-fermentable carbon sources glycerol/ethanol in place of glucose, introducing the low-copy *MRP51* plasmid fully suppressed the Slg^-^ phenotypes displayed by both the *dbp1Δ::hphMX4 ded1-ts* double mutant and *dbp1Δ::hphMX4* single mutant (Figure 1B, row 4 vs. rows 1-2 & row 7 vs 1 and 5). These last findings confirm the conclusion of Powers et al. that the *dbp1Δ::hphMX4* mutation acts in *cis* to reduce expression of chromosomal *MRP51*, reducing cell growth on non-fermentable carbon sources where mitochondrial translation is required for respiration. The *ded1-ts dbp1Δ::hphMX4* double mutant transformed with low-copy *MRP51* shows no Slg^-^ phenotype conferred by *ded1-ts* (Figure 1B, row 7 vs. 1) most likely because the *ded1-ts* product is functional at 30°C (2).

These genetic complementation results demonstrate that the reduced cell growth conferred by *dbp1Δ::hphMX4* in combination with *ded1-ts* on glucose-containing medium results primarily from loss of *DBP1* function rather than reduced *MRP51* expression. Thus, although expression of Dbp1 is low in comparison to Ded1, it makes an appreciable contribution to cell growth when Ded1 function is impaired in cells fermenting glucose. In fact, our ribosome profiling data indicated that *DBP1* expression is up-regulated 2-to 3-fold in *ded1-ts* cells. As such, we consider it likely that the effects of *dbp1Δ::hphMX4* in exacerbating the deleterious effects of the *ded1-ts* mutation on translational efficiencies of many Ded1-hyperdependent mRNAs that we observed on glucose medium reflect loss of Dbp1’s ability to compensate for reduced Ded1 function (6). It remains to be determined whether reduced expression of *MRP51* contributed to the reduced translational efficiencies of any particular mRNAs in *dbp1Δ::hphMX4* or *ded1-ts dbp1Δ::hphMX4* cells.

## ACKNOWLEDGEMENTS

This work was supported by the Intramural Research Program of the National Institutes of Health (FZ & AGH) and by the Department of Biotechnology, India under award number BT/RLF/Re-entry/55/2017 (NDS).

